# Mapping the molecular motions of 5-HT_3_ serotonin-gated channel by Voltage-Clamp Fluorometry

**DOI:** 10.1101/2023.09.28.559960

**Authors:** Laurie Peverini, Sophie Shi, Karima Medjebeur, Pierre-Jean Corringer

## Abstract

The serotonin-gated ion channel (5-HT_3_R) mediates excitatory neuronal communication in the gut and the brain. It is the target for setrons, a class of competitive antagonists widely used as antiemetics, and is involved in several neurological diseases. Cryo-electron microscopy of the 5-HT_3_R in complex with serotonin or setrons revealed that the protein has access to a wide conformational landscape. However, assigning known high-resolution structures to actual states contributing to the physiological response remains a challenge.

In the present study, we used voltage-clamp fluorometry (VCF) to measure simultaneously, for 5-HT_3_R expressed at a cell membrane, conformational changes by fluorescence and channel opening by electrophysiology. Four positions identified by mutational screening report motions around and outside the serotonin-binding site through incorporation of cysteine-tethered rhodamine dyes with or without a nearby quenching tryptophan. VCF recordings show that the 5-HT_3_R has access to four families of conformations endowed with distinct fluorescence signatures: “resting-like” without ligand, “inhibited-like” with setrons, “pre-active-like” with partial agonists and “active-like” (open channel) with partial and strong agonists. Data are remarkably consistent with cryo-EM structures, the fluorescence partners matching respectively Apo, setron-bound, 5-HT bound-closed and 5-HT-bound-open conformations. Data show that strong agonists promote a concerted motion of all fluorescently labelled sensors during activation, while partial agonists, especially when loss-of-function mutations are engineered, stabilize both active and pre-active conformations.

In conclusion, VCF, though the monitoring of electrophysiologically silent conformational changes, illuminates allosteric mechanisms contributing to signal transduction and their differential regulation by important classes of physiological and clinical effectors.

**Significance Statement:** High-resolution structures of serotonin-gated receptors (5-HT_3A_R) have evidenced a wide range of conformations that are challenging to annotate to physiologically relevant states. Voltage-clamp fluorometry allows to investigate the activation of 5-HT_3A_R by simultaneously following molecular motions and electrophysiological states at the plasma membrane. Here, we developed four fluorescent sensors reporting conformational changes at the serotonin binding site and at the extracellular domain and transmembrane domain interface. Investigation of a series of agonists, partial agonists and antagonists show that strong agonists promote a concerted motion of the whole protein during activation, while antagonists and partial agonists stabilize distinct closed-channel conformations. Data offer insights into allosteric mechanisms, unravelling the conformational dynamics of the receptors and helping to annotate high-resolution static structures.

## Introduction

Pentameric ligand-gated ion channels (pLGICs), comprising nicotinic acetylcholine (nAChRs), serotonin (5-HT_3_Rs), glycine (GlyRs) and gamma-amino butyric acid (GABA_A_Rs) receptors are dynamic transmembrane proteins that mediate neuronal and non-neuronal communication (1). Among them, 5-HT_3_Rs are cation-selective excitatory channels expressed in central and peripheral nervous systems, notably in the gut and gut-brain axis (2). 5-HT_3_Rs are encoded by five different genes (*HTR3A*, *HTR3B*, *HTR3C*, *HTR3C*, *HTR3E*), giving rise to five different subunits (A to E) that assemble as homo- or hetero-pentamers. A class of synthetic competitive antagonists called setrons target these receptors and are widely used in the clinic as antiemetics, notably to counteract vomiting and nausea induced by chemotherapy and radiotherapy. 5-HT_3_Rs are also potential targets to treat irritable bowel syndrome and are implicated in neurological diseases such as schizophrenia, Parkinson’s disease, and depression. There is therefore much interest surrounding the molecular mechanisms governing 5-HT_3_Rs function and pharmacological regulation (3–5).

A hallmark of pLGICs, and among them 5-HT_3_R, is that their action is mediated by varying population equilibrium between allosteric states that are differentially stabilized by the binding of ligands (6,7). Agonists promote the allosteric transition between resting (closed channel) and active (open channel) states, while their prolonged application promotes a slower transition to desensitized (agonist-bound closed channel) conformations.

The first high-resolution structure of mouse homo-pentameric 5-HT_3A_R was solved by X-ray crystallography in complex with stabilizing nanobodies, followed by numerous structures solved by Cryo-Electron Microscopy (Cryo-EM) without ligands or in complex with the agonist 5-HT, a series of setron competitive antagonists and an allosteric modulator (8–15). The 5-HT_3A_R displays a typical pLGIC structure, where each subunit consists of a β-sandwich connected with loops composing the extracellular domain (ECD), 4 ⍺-helices composing the transmembrane domain (TMD) and an intracellular domain. Agonists and competitive antagonists bind at the ECD interface between two subunits, at the orthosteric site composed of so-called loops A, B and C from the principal subunit and D, E and F from the complementary subunit, with loop C bridging the subunit interface. In the presence of detergent C12E9 with or without added lipids, or in saposin nanodiscs, solved structures in the absence of orthosteric ligand (Apo) were assigned as resting-like (8,10,15). Those in complex with setrons significantly diverged from the resting-like class and where called inhibited states (9,12–14). The structures captured with bound 5-HT all feature rather similar motions of the ECD, with 5-HT binding promoting a closure (in a capping motion) of loop C around the agonist, a global compaction, and a tilt of each subunit ECD. This generates a marked reorganization of the ECD-TMD interface, including in some cases an outward motion of the M2-M3 loop. However, the structures strongly diverge in the TMD, some featuring an apparently open pore and others a non-conducting pore (9,11). In addition, reconstitution into saposin-based nanodiscs captured distinct open conformations with symmetrical or asymmetrical conformation of the TMD (15). Therefore, these atomic structures led to a complex landscape of more than the three putative conformations (resting, active, desensitized). Furthermore, 5-HT_3_R’s wide conformational landscape accessibility seems to be governed by both orthosteric effectors and the lipids and/or detergents surrounding the TMD. Assigning these high-resolution structures to actual physiological states contributing to the electrophysiological response at the cell membrane remains to be established (16).

Complementary techniques that follow the structural reorganizations of the cell-expressed receptors are thus needed to identify structural motions contributing to gating (i.e., the pathway between resting and activated/open states) and to help annotate high-resolution structures to physiologically-relevant conformational states (16). Voltage-clamp fluorometry (VCF), which monitors receptors expressed at the *Xenopus* oocyte plasma membrane, is well suited for this aim. VCF performs simultaneous measurement of channel opening by two-electrode voltage-clamp electrophysiology (TEVC) and of local protein motions by variation of fluorescence from covalently attached probes (17). This technique has been applied to various pLGICs, notably to human 5-HT_3A_R with the generation of three sensors surrounding the orthosteric site as well as one sensor on the mouse 5-HT_3A_R located at the extracellular entrance of the pore (9,18).

In the present study, we have generated four fluorescent sensors grafted at new locations on the mouse 5-HT_3A_R. We characterized their phenotypes following the binding of various agonists, partial agonists, and clinically relevant antagonists, with or without introduction of loss-of-function mutations. Simultaneous recordings of electrical currents and variations of fluorescence revealed different local and global reorganizations of the ECD depending on the pharmacological conditions. By following the conformational reorganizations leading to both electrophysiologically silent and active conformations, VCF data identify intermediate conformations within or outside the path of 5-HT_3_R activation. These data provide unique structural information on membrane-embedded proteins to annotate known high-resolution structures to physiologically relevant allosteric states.

## Results

### 1) Generation of four fluorescent sensors along the ECD

To guide the design of fluorescent sensors, we inspected the regions undergoing the largest reorganizations in the various Cryo-EM structures, mainly mapping the subunit interface from the apex of the ECD to the ECD-TMD interface and the upper part of the TMD. We introduced cysteines by mutagenesis and covalently labelled them with the fluorescent probe MTS-TAMRA, allowing conjugation of a rhodamine dye with the protein main chain through a flexible 6-atom linker (CH_2_-S-S-CH_2_-NH-CO). Since rhodamine fluorescence is sensitive to its local molecular environment, this generates conformational sensors around the graft position (18). In order to increase the intensity of fluorescent variations and to report more precisely the motion observed, we also implemented the tryptophan-induced quenching method (19) (Figure 1.A.). In this technique, the cysteine-labeled rhodamine is introduced together with a nearby tryptophan since the indole moiety of the Trp side chain robustly quenches rhodamine fluorescence when in Van der Walls contact. Such pairs sense the distance between the fluorophore and indole attachment points and/or local environment, and often amplify the extent of the fluorescence variation when allosteric motions are associated with structural reorganization at this level (39).

**Figure 1.**
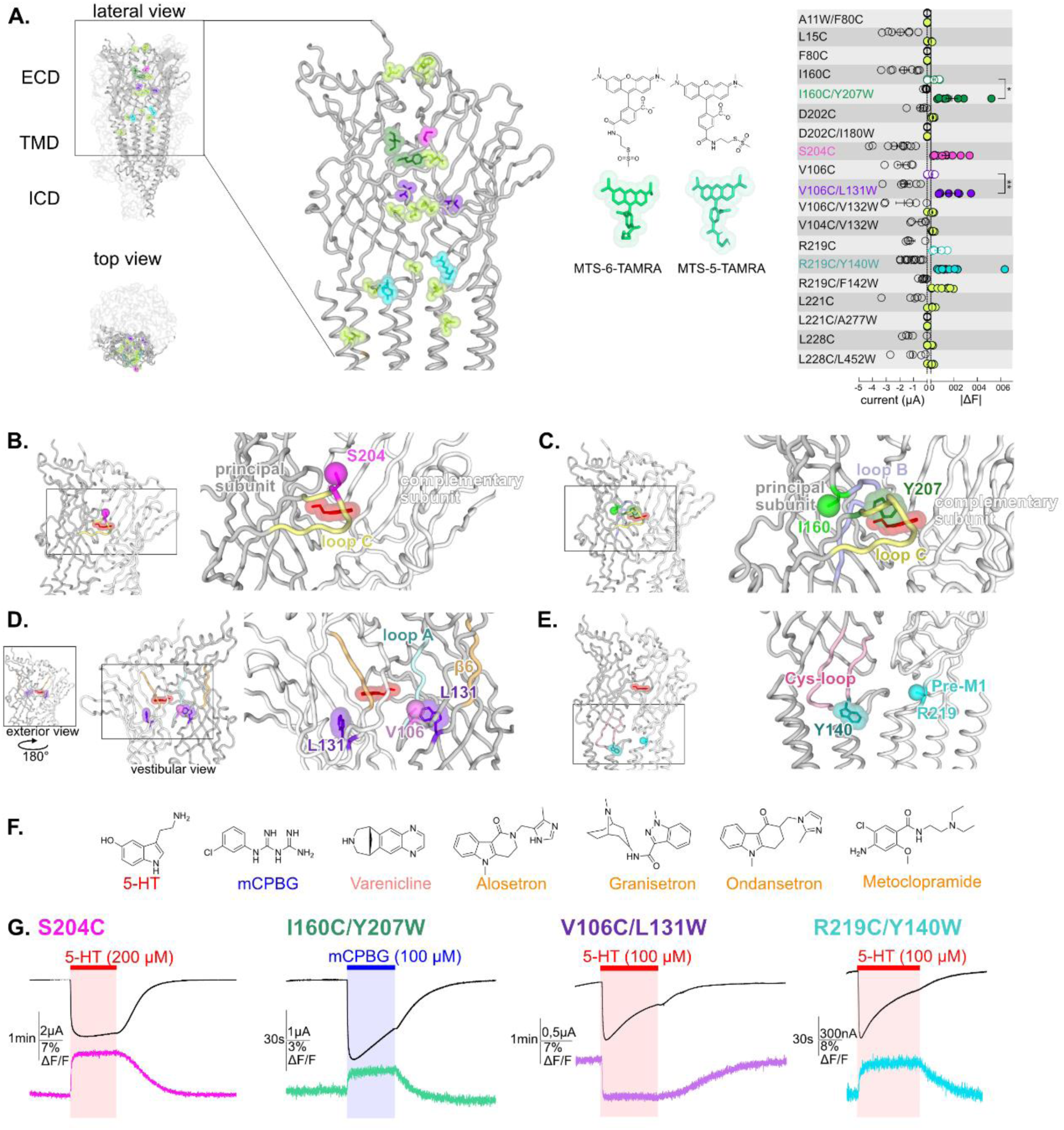
Screening of residues located along the extracellular domain (ECD) of m5-HT3AR as potential anchors for fluorescent probes and establishment of four reporter sensors. **A.** Left panel: visualization of the residues mutated, TAMRA-labelled and tested for variation of current and fluorescence on the Apo Cryo-EM structure of m5-HT_3A_ (from Basak et al., 2018), in lateral (zoom on the ECD) and top view with two subunits in cartoon representation and the three others represented in surface mode (PDB: 6BE1). Middle panel: representation of the two isomers of the labeling fluorescent probe mixture used in this study, MTS-TAMRA (5(6)-carboxytetramethylrhodamine methanethiosulfonate). Right panel: representation of currents evoked by 50-100 µM of 5-HT on the mutants and of the absolute values of variation of fluorescence (difference between the baseline fluorescence and steady-state fluorescence upon 5-HT perfusion) recorded simultaneously. Note the representation of the four selected sensors: in dark green, I160C/Y207W; in magenta, S204C; in purple, V106C/L131W and in cyan, R219C/Y140W. **B.** The sensor S204C, shown on 5-HT bound conformation (5-HT represented in red; PDB: 6DG8), is located on loop C. Note that the fluorescence variation is significatively increased by the addition of the tryptophan in the sensors I160C/Y207W and V106C/L131W. (unpaired t-tests, I160C/Y207W versus I160C P value = 0,013 (*); V106C/L131W versus V106C P value = 0,0081 (**)) **C.** The sensor I160C/Y207W where I160C is the point of labelling with MTS-TAMRA and is represented in ball representation in fluorescent green, Y207 is mutated into tryptophan and colored in dark green. **D.** The sensor V106C/L131W where V106C is represented in ball representation in purple and L131 is mutated into tryptophan and colored in purple. Note the vestibular positioning of this sensor. **E.** The sensor R219C/Y140W, where R219C, located in pre-M1 loop, is represented in ball representation in cyan and Y140, located in the Cys-loop, is mutated into tryptophan, and colored in dark cyan. **F.** Molecular structures of the ligands used in this study: agonists (in red, 5-HT; in blue, mCPBG; in salmon, varenicline) and antagonists (all represented in orange, A: alosetron, G: granisetron, O: ondansetron, M: metoclopramide). **G.** Effect of desensitization on the dynamic of the fluorescence recordings. Examples of desensitizing currents promoted by prolonged perfusion of strong agonists (to elicit robust desensitization): mCPBG perfused on the sensor I160C/Y207W or 5-HT perfused on the sensors S204C, V106C/L131W and R219C/Y140W. Traces show that the fluorescent signal remains stable during desensitization for all the sensors. Note that the differences in the desensitization kinetics of the displayed traces are due to the variability of different oocyte batches, since the four conditions do not show significant differences in desensitization kinetics (Fig S2).

Nineteen TAMRA-labelled cysteine mutants were screened by VCF under perfusion of high concentration of 5-HT (Figure 1. A). We verified that the mouse wild-type 5-HT_3_R (m5-HT_3_R), that does not carry any single cysteine within the ECD, yielded robust 5-HT-elicited currents upon treatment with MTS-TAMRA but no changes in fluorescence intensity. The screening allowed for the selection of four mutants (one cysteine and three cysteine/tryptophan pairs) endowed with robust variations of both current and fluorescence.

We first identified the single mutation S204C, which is located near the tip of loop C, facing the adjacent subunit and the ECD interface above the orthosteric site (Fig 1.B). Second, three fluorophore/quencher (cysteine/tryptophan) pairs were selected. In each case, the measured 5-HT-elicited variation of fluorescence (ΔF) was much smaller or unmeasurable when the cysteine but not the tryptophan was introduced, indicating that the dequenching or quenching observed is dominated by Van der Walls interaction between the rhodamine dye and the indole of the tryptophan (Figure S1.C.). The pairs are the following:

1) I160C/Y207W: I160C is located in close proximity to the orthosteric site but outside the pocket behind the base of loop C, while Y207W lies inside the pocket nearby bound ligands in agonist and antagonist co-structures. Of note, the aromatic side chain of Y207 contributes to the binding and gating in 5-HT_3_R, with a strong effect of alanine and serine mutations, but comparatively weaker effect of phenylalanine or unnatural aromatic mutations (20). The pair is located away from the subunit interface thus reporting tertiary reorganizations (Fig 1.C).
2) V106C/L131W: V106C is facing the vestibule of the ECD, a large water-accessible channel with several constrictions and ring of charges that lies above the TMD ion channel. L131W is positioned more profoundly in each ECD, its side chain remaining visually accessible through the vestibule. Given the proximity of all subunits in this confined space, the observed fluorescence variation can potentially arise from quenching by tryptophan of any of the five subunits (Fig 1.D). Of note, we verified that the observed fluorescent quenching is not caused by direct collisional quenching with the hydroxy indole group of perfused 5-HT, that arises at concentrations above 200 μM (see an example of direct quenching by 5-HT perfused at 1000 µM, Figure S1.C.).
3) R219C/Y140W are positioned at the ECD/TMD interface. R219C is located in the pre-M1 area and Y140W in the highly conserved Cys-loop. Both positions flank the M2-M3 loop that undergoes a major outward motion in some agonist-bound structures (Fig 1.E).

### 2) The four sensors monitor fast motions not related to desensitization

The four sensors were recorded on a custom VCF chamber where the ligands are perfused only on the portion of the oocyte from which the fluorescence emission is collected (21). This ensures that the same population of receptors is recorded in current and fluorescence simultaneously. For all sensors, perfusion of a high concentration of strong agonists (5-HT or mCPBG, selected depending on the particular sensor, see dedicated section to each sensor) elicit currents reaching maximal value in a few seconds, followed by desensitization appearing with much slower kinetics (Fig 1.G). To measure the desensitization kinetics, we performed parallel measurements on a dedicated TEVC setup equipped with a fast perfusion system allowing solution exchange in less than a hundred milliseconds (22). This shows that all sensors display comparable desensitization kinetics to that of the wild-type receptor (WT), evaluated in the 50-150 s^−1^ range though mono-exponential fitting (Fig S2). Thus, activation and desensitization appear well separated in time in the VCF setup. This allows a reasonable evaluation of the extent of activation by measuring the amplitude of the peak current. Concentration-response curves measured at this peak current show that, for all sensors, labelling by MTS-TAMRA has no significant effect in terms of EC_50_ current (EC_50_c) (Fig S1.B).

In fluorescence, labelled S204C, I160C/Y207W and R219C/Y140W show robust agonist-elicited dequenching and V106C/L131W agonist-elicited quenching. In all cases, the rise-time of the fluorescence variations (ΔFs) are in the same range to that of the rise-time of the currents (1-10 s range depending on the particular sensor and the agonist concentration, Table S1). In contrast, prolonged applications of agonist show no significant variation of the fluorescence during the desensitization phase (with the fluorescent signal remaining stable), providing evidence that the sensors do not report movements related to desensitization (Fig 1.G). As far as the four sensors are concerned, VCF data suggest that the ECD (and its labels) does not undergo notable conformational changes between activated and desensitized states.

### 3) Competitive antagonists elicit agonist-like reorganizations at the orthosteric site that do not spread to the lower part of the ECD

We first explored the action of competitive antagonists on the various sensors. To this end, we applied to the same oocyte saturating concentrations of the agonists 5-HT, mCPBG (*m*-chlorophenylbiguanide) and var (varenicline) as well as a selection of four competitive antagonists of different molecular structure: alosetron, granisetron, ondansetron and metoclopramide (23–25). We used a 3 µM concentration of each antagonist, which is far above their nanomolar binding affinities measured on the WT receptor.

On the two sensors neighboring the orthosteric site, antagonists produce no current, but robust ΔFs are observed in the same direction as agonists do (Fig 2.A and 2.C). In addition, the amplitude of ΔFs evoked by antagonists are in the same range than that of agonists for granisetron and ondansetron while it is significantly higher for alosetron and lower for metoclopramide. This suggests that, locally, antagonists elicit similar reorganizations as agonists do but with various amplitudes (Figure 2.A. and 2.C. left panels).

**Figure 2.**
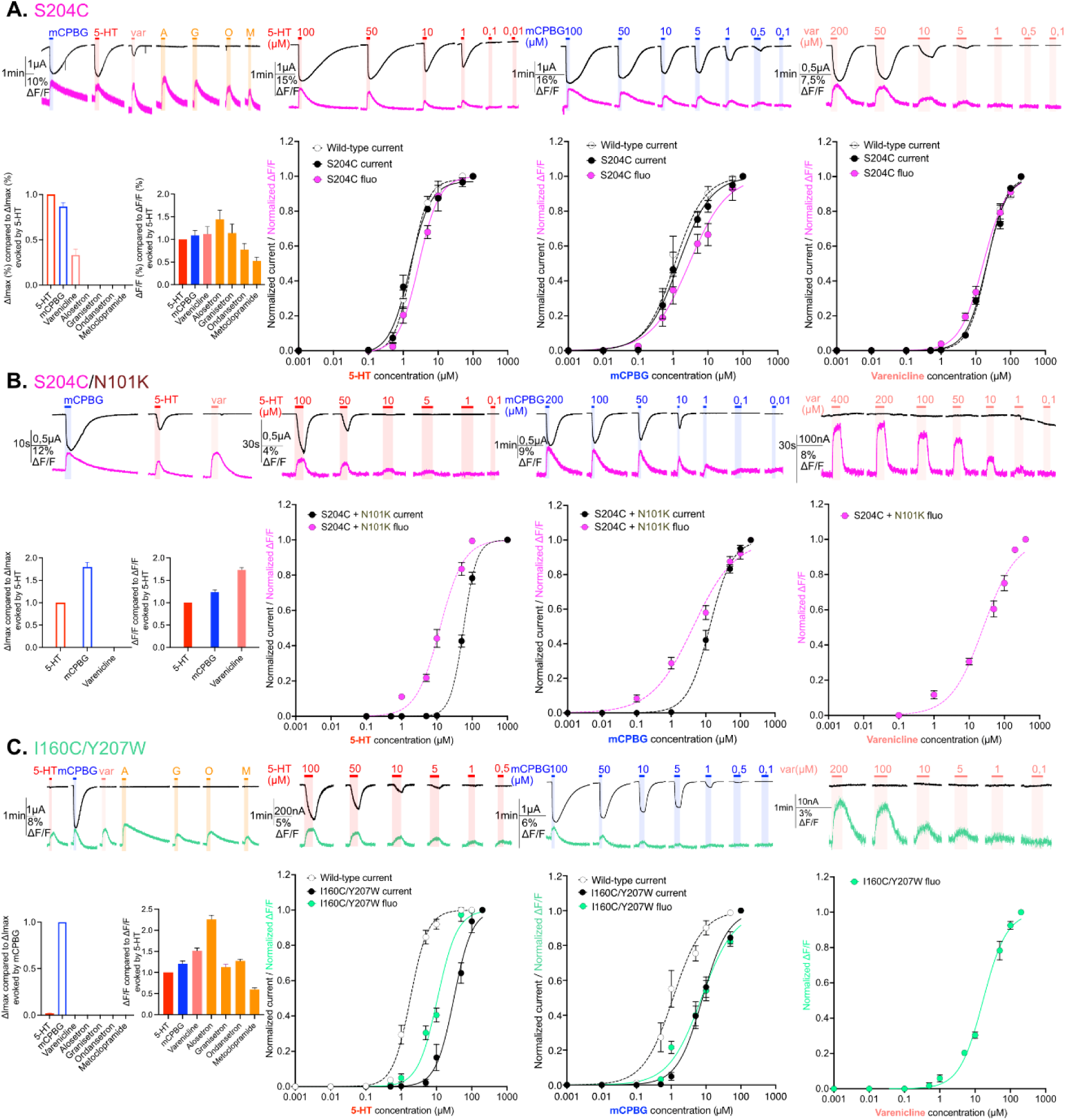
Electrophysiological and fluorescence characterization of S204C and I160C/Y207W, two sensors located in proximity to the ligand-binding site. **A. Exploration and characterization of the sensor S204C.** Left upper panel: macroscopic ligand-gated currents (in black) and fluorescence (in magenta) recorded at −60mV on S204C labelled with MTS-TAMRA evoked by saturating concentrations of agonists (in red, 5-HT, 200µM ; in blue, mCPBG, 200µM; in salmon, varenicline, 400µM) and antagonists (all represented in orange, A: alosetron, G: granisetron, O: ondansetron, M: metoclopramide, all at 3µM). Left bottom panel: graphical representation of ligand-induced relative changes of current and fluorescence compared to 5-HT. The normalized values for all the ligands are compared with the mean values obtained for 5-HT. Middle left top panel: representative recording of current and fluorescence variations of S204C labelled with MTS-TAMRA upon different concentrations of perfused 5-HT. Middle left bottom panel: dose-response curves for ΔI (black) and ΔF (magenta) with mean and SEM (normalized by the maximum current of each oocytes) for application of 5-HT. Middle right top panel: representative recording of current and fluorescence variations of S204C labelled with MTS-TAMRA upon different concentrations of perfused mCPBG. Middle right bottom panel: dose-response curves for ΔI (black) and ΔF (magenta) for application of mCPBG. Right top panel: representative recording of current and fluorescence variations of S204C labelled with MTS-TAMRA upon different concentrations of perfused varenicline. Right bottom panel: dose-response curves for ΔI (black) and ΔF (magenta) for application of varenicline. **B. Effect of loss-of-function mutation N101K on the sensor S204C.** Same experiments and legends as for panel A. but the construct here is the sensor S204C with the additional loss-of-function mutation N101K. **C. Exploration and characterization of the sensor I160C/Y207W.** Same experiments and legends as for panel A, but here currents are represented in black and fluorescence in green. Note also that for the left lower panel, currents and fluorescence have been compared to mCPBG instead of 5-HT.

In contrast, the sensors at the vestibular site (V106C/L131W) and at the ECD-TMD interface (R219C/Y140W) show no antagonist-elicited current nor ΔF, suggesting that the conformational effects elicited by these competitive antagonists do not spread to the vestibule and lower part of the ECD (Fig 3.A. and 3.C, left panels).

**Figure 3.**
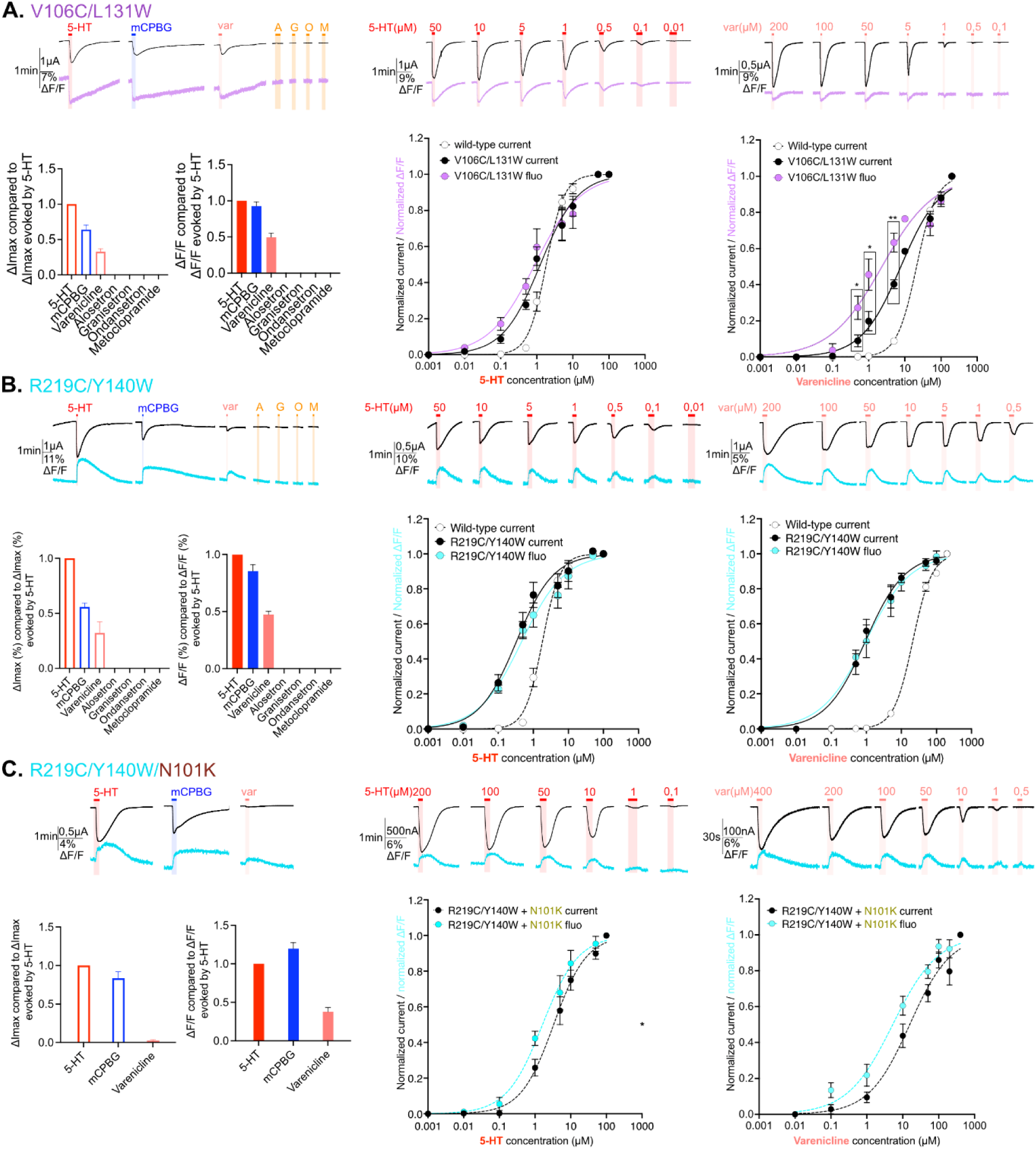
Electrophysiological and fluorescence characterization of V106C/L131W, located in the extracellular vestibular area and R219C/Y140W positioned at the interface area between ECD and TMD. **A. Exploration and characterization of the sensor V106C/L131W**. Left top panel: macroscopic ligand-gated currents (in black) and fluorescence (in purple) recorded at −60mV on the construct with the sensor V106C/L131W labelled with MTS-TAMRA evoked by saturating concentrations of agonists (in red, 5-HT; in blue, mCPBG; in salmon, varenicline) and antagonists (all represented in orange, A: alosetron, G: granisetron, O: ondansetron, M: metoclopramide). Left bottom panel: graphical representation of ligand-induced relative changes of current and fluorescence compared to 5-HT. The normalized values for all the ligands are compared with the mean values obtained for 5-HT. Middle top panel: representative recording of current and fluorescence variations of V106C/L131W labelled with MTS-TAMRA upon different concentrations of perfused 5-HT. Middle bottom panel: dose-response curves for ΔI (black) and ΔF (purple) with mean and SEM (normalized by the maximum current of each oocyte) for application of 5-HT. Right top panel: representative recording of current and fluorescence variations of V106C/L131W labelled with MTS-TAMRA upon different concentrations of perfused varenicline. Right bottom panel: dose-response curves for ΔI (black) and ΔF (magenta) with mean and SEM (normalized by the maximum current of each oocyte) for application of varenicline. Note the significative difference between dose response of current and fluorescence at 0.5, 1 and 5 µM of perfused varenicline (unpaired t-test). **B. Exploration and characterization of the sensor R219C/Y140W.** Same experiments and legends as for panel A. but the construct here is the sensor R219C/Y140W, current is represented in black and fluorescence in cyan (in trace recordings and dose-response representations). **C. Effect of loss-of-function mutation N101K on the sensor R219C/Y140W.** Same experiments and legends as for the panel A. but the construct here is the sensor R219C/Y140W with the additional loss-of-function mutation N101K, current is represented in black and fluorescence in cyan (in trace recordings and dose-response representations).

### 4) Correlation between the orthosteric site motions and ion channel activation

The sensor S204C displays electrophysiological properties identical to that of the WT, both in terms of agonist potency and efficacy. EC_50_c are identical between S204C and WT, i.e. 1.8 and 1.8 µM for 5-HT, 1.4 and 1.1 µM for mCPBG, and 20.6 and 21.1 µM for var, respectively (Table 2). mCPBG appears as a near full agonist, eliciting 89% and 92% of maximal 5-HT currents on S204C and WT, respectively, while var appears as a partial agonist eliciting only 33% and 39% of maximal 5-HT currents on S204C and WT, respectively (Table 1). All data on WT are consistent with TEVC data reported in the literature (24).

**Table 1:** Current and fluorescence maximum evoked by ligands (agonist and antagonists) on m5-HT_3A_ mutants.

| Construct | Molecule | $\Delta I_{\max}$ molecule compared to $\Delta I_{\max}^{5-HT}$ +/- SEM | $\Delta F_{\max}$ molecule compared to $\Delta F_{\max}^{5-HT}$ +/- SEM | n |
| --- | --- | --- | --- | --- |
| **Wild-type** | mCPBG | 0.92 +/- 0.03 | / | 6 |
|  | Varenicline | 0.39 +/- 0.06 | / |  |
|  | aloseptron | 0 | / |  |
|  | granisetron | 0 | / |  |
|  | ondasetron | 0 | / |  |
|  | metoclopramide | 0 | / |  |
| **S204C + MTS-TAMRA** | mCPBG | 0.89 +/- 0.04 | 1.09 +/- 0.105 | 6 |
|  | Varenicline | 0.33 +/- 0.06 | 1.12 +/- 0.16 |  |
|  | aloseptron | 0 | 1.44 +/- 0.20 |  |
|  | granisetron | 0 | 1.14 +/- 0.19 |  |
|  | ondasetron | 0 | 0.78 +/- 0.13 |  |
|  | metoclopramide | 0 | 0.52 +/- 0.08 |  |
| **I160C/Y207W + MTS-TAMRA** | mCPBG | *5-HT – 0.02 +/- 0.004 | 1.21 +/- 0.06 | 8 |
|  | Varenicline | *0 | 1.52 +/- 0.06 |  |
|  | aloseptron | *0 | 2.26 +/- 0.10 |  |
|  | granisetron | *0 | 1.12 +/- 0.07 |  |
|  | ondasetron | *0 | 1.27 +/- 0.03 |  |
|  | metoclopramide | *0 | 0.60 +/- 0.04 |  |
| **V106C/L131W + MTS-TAMRA** | mCPBG | 0.64 +/- 0.06 | 0.93 +/- 0.06 | 6 |
|  | Varenicline | 0.33 +/- 0.04 | 0.49 +/- 0.06 |  |
|  | aloseptron | 0 | 0 |  |
|  | granisetron | 0 | 0 |  |
|  | ondasetron | 0 | 0 |  |
|  | metoclopramide | 0 | 0 |  |
| **R219C/Y140W + MTS-TAMRA** | mCPBG | 0.62 +/- 0.07 | 0.94 +/- 0.09 | 6 |
|  | Varenicline | 0.34 +/- 0.08 | 0.47 +/- 0.02 |  |
|  | aloseptron | 0 | 0 |  |
|  | granisetron | 0 | 0 |  |
|  | ondasetron | 0 | 0 |  |
|  | metoclopramide | 0 | 0 |  |
| **S204C + N101K - MTS-TAMRA** | mCPBG | 1.80 +/- 0.11 | 1.23 +/- 0.05 | 8 |
|  | Varenicline | 0 | 1.73 +/- 0.05 |  |
| **R219C/Y140W + N101K - MTS-TAMRA** | mCPBG | 0.83 +/- 0.08 | 1.20 +/- 0.08 | 7 |
|  | Varenicline | 0.03 +/- 0.01 | 0.38 +/- 0.06 |  |

**Table 2:** EC_50_ values for current (EC_50_c) and fluorescence (EC_50_f) responses to agonists (5-HT, mCPBG, varenicline) at labeled and unlabeled m5-HT_3A_ mutants. The top part of the table represents the characterization of the sensors with different ligands and the associated controls (rows 1 to 13) and the second part the additional allosteric mutations added (rows 14 to 16).

| Construct | Molecule | EC<sub>50c</sub> (μM) +/- SEM | n<sub>Hillc</sub> +/- SEM | EC<sub>50f</sub> (μM) +/- SEM | n<sub>Hillf</sub> +/- SEM | n |
| --- | --- | --- | --- | --- | --- | --- |
| Wild-type (WT) | 5-HT | 1.83 +/- 0.12 | 1.74 +/- 0.13 | / | / | 8 |
|  | mCPBG | 1.13 +/- 0.15 | 1.02 +/- 0.14 | / | / | 6 |
|  | Varenicline | 21.09 +/- 1.03 | 1.595 +/- 0.07 | / | / | 5 |
| WT + MTS-TAMRA | 5-HT | 3.43 +/- 0.24 | 1.98 +/- 0.12 | / | / | 5 |
| I160C/Y207W + MTS-TAMRA | 5-HT | 31.22 +/- 2.76 | 1.69 +/- 0.18 | 10.73 +/- 0.62 | 1.56 +/- 0.14 | 7 |
|  | mCPBG | 7.99 +/- 0.58 | 1.19 +/- 0.105 | 7.46 +/- 0.755 | 0.92 +/- 0.08 | 8 |
|  | Varenicline | No current | / | 16.80 +/- 0.76 | 1.29 +/- 0.06 | 7 |
| I160C/Y207W unlabelled | 5-HT | 47.33 +/- 4.72 | 1.19 +/- 0.135 | / | / | 6 |
|  | mCPBG | 2.73 +/- 0.17 | 1.59 +/- 0.11 | / | / | 8 |
|  | Varenicline | No current | / | / | / | / |
| I160C + MTS-TAMRA | 5-HT | 31.01 +/- 1.64 | 1.73 +/- 0.11 | / | / | 5 |
| S204C + MTS-TAMRA | 5-HT | 1.78 +/- 0.13 | 1.39 +/- 0.11 | 2.86 +/- 0.248 | 1.51 +/- 0.14 | 6 |
|  | mCPBG | 1.45 +/- 0.18 | 0.94 +/- 0.10 | 2.90 +/- 0.40 | 0.80 +/- 0.08 | 6 |
|  | Varenicline | 20.67 +/- 0.705 | 1.48 +/- 0.05 | 16.31 +/- 0.78 | 1.29 +/- 0.06 | 7 |
| S204C unlabelled | 5-HT | 1.93 +/- 0.25 | 1.08 +/- 0.12 | / | / | 6 |
|  | mCPBG | 0.54 +/- 0.02 | 1.94 +/- 0.16 | / | / | 6 |
|  | Varenicline | 17.105 +/- 1.12 | 1.385 +/- 0.07 | / | / | 7 |
| V106C/L131W + MTS-TAMRA | 5-HT | 1.25 +/- 0.155 | 0.84 +/- 0.08 | 0.88 +/- 0.16 | 0.68 +/- 0.08 | 5 |
|  | Varenicline | 7.745 +/- 0.775 | 0.80 +/- 0.05 | 2.33 +/- 0.45 | 0.60 +/- 0.06 | 5 |
| V106C/L131W unlabelled | 5-HT | 1.91 +/- 0.17 | 0.99 +/- 0.07 | / | / | 9 |
|  | Varenicline | 6.06 +/- 0.32 | 1.47 +/- 0.12 | / | / | 5 |
| V106C + MTS-TAMRA | 5-HT | 4.47 +/- 0.55 | 1.08 +/- 0.13 | / | / | 5 |
| R219C/Y140W + MTS-TAMRA | 5-HT | 0.33 +/- 0.05 | 0.82 +/- 0.10 | 0.47 +/- 0.07 | 0.71 +/- 0.07 | 8 |
|  | Varenicline | 0.94 +/- 0.12 | 0.77 +/- 0.08 | 0.96 +/- 0.19 | 0.68 +/- 0.10 | 6 |
| R219C/Y140W unlabelled | 5-HT | 0.60 +/- 0.01 | 0.94 +/- 0.10 | / | / | 7 |
|  | Varenicline | 1.575 +/- 0.28 | 0.875 +/- 0.12 | / | / | 6 |
| R219C + MTS-TAMRA | 5-HT | 0.11 +/- 0.03 | 0.92 +/- 0.21 | 0.185 +/- 0.04 | 0.94 +/- 0.15 | 6 |
| WT + N101K | 5-HT | 66.67 +/- 3.11 | 1.95 +/- 0.185 | / | / | 6 |
| S204C + N101K | 5-HT | 57.11 +/- 1.49 | 2.33 +/- 0.17 | 12.19 +/- 0.90 | 1.285 +/- 0.11 | 6 |
|  | varenicline | No current | / | 25.30 +/- 2.46 | 0.94 +/- 0.07 | 7 |
|  | mCPBG | 13.10 +/- 0.79 | 1.345 +/- 0.10 | 4.365 +/- 0.495 | 0.72 +/- 0.05 | 7 |
| R219C/Y140W + N101K | 5-HT | 3.32 +/- 0.39 | 0.94 +/- 0.10 | 1.62 +/- 0.27 | 0.86 +/- 0.11 | 9 |
|  | varenicline | 15.65 +/- 2.70 | 0.76 +/- 0.09 | 5.25 +/- 0.95 | 0.69 +/- 0.07 | 6 |

The strong agonists 5-HT and mCPBG elicit similar maximum fluorescence variation (ΔFmax) at saturation. Concentration-response curves show that the variations of current (ΔIs) and fluorescence are well correlated with similar EC_50_ fluorescence (EC_50_f) and EC_50_c. This shows that agonists promote local reorganizations around the binding site that are correlated to the opening of the channel, suggesting a concerted motion of the orthosteric site and the ion channel during activation. As for strong agonists, var displays similar EC_50_f and EC_50_c. However, it displays a ΔFmax similar to that of strong agonists despite activating only 33% of the current. Therefore, var at saturation promotes full reorganization of the binding site suggesting that among the population displaying a variation of fluorescence, only a fraction shows an open channel (Fig 2.A).

To further investigate the S204C sensor on a different allosteric background, we introduced the strong loss-of function mutation N101K. Substitution of N101 by several natural or unnatural amino acids indeed produces a marked increase in agonist EC_50_ (26,27). N101 is located just below the orthosteric site on loop A and has been proposed to contribute to the allosteric network coupling the orthosteric site to the ion channel. In combination with S204C, N101K displays, as expected, a loss-of-function phenotype characterized by: 1/ a 31-fold and 9-fold increase in EC_50_c for 5-HT and mCPBG, respectively, 2/ a decreasing efficacy of 5-HT as compared to mCPBG, as already reported and 3/ a decreasing efficacy of var yielding no detectable current (Fig 2.B and Tables 1 and 2). Interestingly, the effects on the fluorescence variations were comparatively weaker, N101K causing only a 4-fold and a 1.4-fold increase in EC_50_f for 5-HT and mCPBG, respectively, while var causes a ΔF higher than 5-HT at saturation (173%, figure 2.B. left panel and Table 1) with a 1.6-fold increase in EC_50_f (Table 2 and Figure 2.B.). Therefore, N101K on S204C decorrelates the ΔF and ΔI. For 5-HT and mCPBG, ΔFs are already robust at agonists concentrations not eliciting sizable currents (for instance at 5 µM 5-HT and 1 µM mCPBG, see Figure 2.B.). For var, it elicits quantitative ΔF but no current.

Finally, the sensor I160C/Y207W displays by itself a marked loss-of-function phenotype characterized by: 1/ a 7-fold and 17-fold increase in EC_50_c for mCPBG and 5-HT as compared to the WT, respectively (Table 2) and 2/ at saturation, mCPBG is the most efficient agonist, while 5-HT elicits only 2 % of its currents and var fails to activate the channel (Table 1, Fig 2.C. left panel). Of note, this phenotype is caused by both I160C and Y207W mutations, that individually cause a 17-fold and 12-fold increase in the EC_50_c for 5-HT, respectively (Fig S1.A. right panel). Given the position of these mutations within and around the orthosteric site, their effect is likely due to a mix of alteration of both gating and binding affinity of the agonists. The phenotype of I160C/Y207W resembles that of S204C/N101K, with a leftward shift of the ΔF over the ΔI curve for the partial agonist 5-HT, and robust ΔF but no ΔI for var (Fig 2.C).

Altogether, VCF data obtained for these two sensors close to the binding site highlight three families of conformations, the apo conformations, the agonist-bound active conformations characterized by a concerted reorganization of the orthosteric site and the ion channel (opening), and ligand-bound intermediate conformations characterized by reorganization of the orthosteric site with a closed channel. On the S204C that displays a WT-like phenotype, these intermediate conformations are partially populated for the partial agonist var. On the two loss-of-function backgrounds (S204C/N101K and I160C/Y207W), these intermediates are fully stabilized by var that behaves as an antagonist and are partially populated for 5-HT and mCPBG at sub-saturating concentrations.

### 5) Correlation between vestibular and ECD-TMD interface motions and ion channel activation

VCF recordings of both V106C/L131W and R219C/Y140W sensors show reproducible currents upon mCPBG perfusion at saturation but not in the case of long recordings required for dose-response curves, where they were non-reproducible. In contrast, 5-HT and var elicit robust and reproducible currents. For both sensors, quantification of the desensitization kinetics on the fast-perfusion TEVC setup suggest a slight, although non-significant, increase in desensitization kinetics for 5-HT and var but shows surprisingly very fast desensitization kinetics for mCPBG (Fig S2). This explains why, in the VCF setup endowed with a relatively slow perfusion system, the very transient mCPBG-elicited activation peak is truncated by desensitization, precluding its further analysis.

V106C/L131W and R219C/Y140W both display a gain-of-function phenotype that is moderate for the former (1.5-fold and 2.7-fold decrease EC_50_c for 5-HT and var, respectively), and more pronounced for the latter (5.5-fold and 22.5-fold decrease EC_50_c for 5-HT and var, respectively). For both sensors, var appears as a partial agonist eliciting around 30% of the 5-HT currents (Fig 3.A, 3.C and Table 1). We also combined R219C/Y140W with the loss-of function N101K mutation. N101K reverts the R219C/Y140W gain-of-function phenotype, the triple mutant R219C/Y140W/N101K displaying EC_50_c of 5-HT and var close to that of the WT (Fig 3.B).

These three sensors show a good correlation between EC_50_c and EC_50_f for the strong agonist 5-HT, suggesting that the motions reported by fluorescence at the vestibule and ECD-TMD interface are concerted with channel opening. For the partial agonist var on V106C/L131W and R219C/Y140W, the ΔFmax (40-50% of that of 5-HT) is comparable to the ΔImax (30-40% of that of 5-HT), suggesting in the first approximation that var at saturation promotes the same transition as 5-HT, but only partially. However, concentration-response curves on V106C/L131W show a small yet significant decorrelation of fluorescence and current, suggesting the occurrence of intermediate conformations. R219C/Y140W/N101K confirms this idea with var. On this construct, var acts as a strongly partial agonist eliciting at saturation only 3% of the 5-HT currents, while it displays 37% of the ΔF evoked by 5-HT (Fig 3.B. left panel). In addition, var shows a leftward shift of the fluorescence dose-response curve as compared to the current dose response curve (with EC_50_c being 15.6 µM and EC_50_f is 5.2 µM) (Fig 2.B, right panel).

In conclusion, as for orthosteric-located sensors, fluorescent and current signals are well correlated for strong agonists, pointing to a concerted mechanism. In contrast, the partial agonist var on V106C/L131W and especially on R219C/Y140W/N101K shows a decorrelation of fluorescence and current, pointing to the contribution of distinct states endowed with robust ΔF but no ΔI.

## Discussion

In this work, we undertook a screening of 19 mutants of the m5-HT_3A_R with an aim to identify fluorescent sensors within the ECD. Around the orthosteric site, we find that cysteine incorporation often yields ΔF suitable for VCF investigation. In contrast, outside the orthosteric site all single-cysteine mutants investigated failed. This observation parallels a previous VCF screening on the h5-HT_3A_R that identified four suitable sensors around the orthosteric site but none among the seven additional positions tested in the pre-M1 region (18). Of note, we have previously investigated a sensor on the top of M2 helices on m5-HT_3A_R (9). In the pLGIC family, the ⍺1-GlyR has also been extensively studied by VCF and many positions were screened for cysteine incorporation (21,28–31). In contrast to the 5-HT_3_R, most positions show robust ΔF outside the orthosteric site. This suggests that the ⍺1-GlyR undergoes gating reorganizations of larger amplitude than that of the 5-HT_3_R. Still, we succeeded in designing two sensors at the vestibule and the ECD-TMD interface by implementing the tryptophan-induced quenching method, further illustrating that this technique is suitable to monitor subtle conformational changes when the fluorophore-quencher pair is properly engineered.

The mapping of the 5-HT_3_R has enabled us to generate four fluorescence sensors of the protein conformations. They are located at three distant positions within the ECD 3D structure, thereby constituting reference points to infer its global conformation. Simultaneous steady-state concentration-response relationships for the fluorescence and current responses reveal important ligand-elicited protein motions and their relationship with activation.

We first show that agonists and antagonists elicit similar local reorganizations around orthosteric sensors. For both S204C and I160C/Y207W, all tested ligands at saturating concentrations elicit fluorescent dequenching signals. All agonists, partial agonists, granisetron and ondansetron elicit ΔF of similar amplitude, while alosetron compared to metoclopramide display significantly higher and lower ΔF, respectively. These results are in remarkable concordance with Cryo-EM structures. They show that both 5-HT and a series of setron antagonists promote locally a similar reorganization of the orthosteric site, with an inward displacement of the landmark loop C that caps the bound ligand (Fig S3) (9,11). Furthermore, subjecting these structures to molecular dynamic simulations suggests that different antagonists promote different degrees of local conformational changes (12,13). Among setrons, alosetron is the one promoting the largest conformational effects, consistent with VCF data. Of note, the ligand-elicited dequenching signal of the I160C/Y207W is also consistent with the Cryo-EM structures, since the capping of loop C is predicted to reduce the accessibility of the indole side chain to the rhodamine dye grafted on the other side of the loop. Interestingly, three other fluorescence sensors around the orthosteric site were already reported on the human 5-HT_3A_R on loop C (M223C), D (Y89C) and E (Q146C). They also report that agonists and antagonists equally elicit fluorescence dequenching, with ΔF intensities strongly ligand-dependent for loop C and E, but not for loop D (18).

We also show that antagonist-elicited reorganizations do not spread to the vestibular and ECD-TMD sensors. Indeed, the sensors at the ECD-TMD interface (R219C/Y140W) and at the middle vestibule (V106C/L131W) show robust ΔF upon perfusion of agonists, but no effect for antagonists. Cryo-EM structures show no setron-elicited change at the ECD-TMD interface, especially no motion of the M2-M3 loop that lies in between the attachment points of the fluorophore and the indole of the tryptophan in R219C/Y140W (Fig S3) (9,13,14). Likewise, setrons elicit small reorganizations of the architecture of the vestibule (measured as a slightly increased radius of the constriction at K108 and D105), while 5-HT elicits stronger reorganizations at this level. Overall, VCF data are in excellent agreement with Cryo-EM data, providing evidence that high resolution structures of setron-inhibited states represent pertinent snapshots of actual allosteric states at the plasma membrane.

Steady-state concentration-response relationships support that strong agonists elicit an apparently concerted reorganization of the global protein structure. Indeed, for most constructs, those agonists eliciting the maximal response currents amongst all tested agonists show no difference in ΔF and ΔI concentration-response curves. This is the case for 5-HT, that maximally activates the S204C, V106C/L131W and R219C/Y140W sensors and for mCPBG that maximally activates the S204C and I160C/Y207W sensors. This suggests a concerted mechanism where local motions around the four sensors happen simultaneously with the pore opening. These data are thus accounted by a simple two-state model, where the receptor is in equilibrium between a resting and an active state. VCF measurements are also remarkably coherent with the 5-HT-bound structures showing an open pore: PDB 6HIN (called F), PDB 6DG8 (called State 2) and PDB Y5A (called serotonin-bound state). Indeed, all of them show symmetrical and global reorganizations of the ECD involving loop C capping, vestibular reorganization and M2-M3 reorganization (Fig S3) (9,11,15).

However, we found that partial agonists stabilize additional intermediate conformations characterized by fluorescence variations but no current, especially when loss-of-function mutations are engineered. Intermediate conformations are observed for the two sensors outside the orthosteric site. Indeed, for R219C/Y140W sensor in combination with N101K, var elicits at saturation only 3% of 5-HT currents but promotes a robust ΔF signal corresponding to 38% of that of 5-HT. In addition, the ΔF concentration-response curve appears slightly left-shifted as compared to the ΔI curve, indicating that the ΔF/ΔI ratio is even higher at low var concentrations. These data clearly show that var stabilizes in majority intermediate conformations where the ECD-TMD interface has moved, generating a ΔF, but where the channel remains closed. A similar although milder “intermediate” phenotype is seen for var on the V106C/L131W sensor.

Within the gating pathway, these intermediate conformations may *a priori* contribute either to the activation transition (pre-active state) or to the desensitization transition (fast or slow desensitized states). However, VCF data strongly argue for the former hypothesis. Indeed, for all experiments including var partial activation of V106C/L131W, the rise-times of ΔF are in the same range to that of ΔI, desensitization being much slower and associated with no significant ΔF. This shows that the intermediate states are recruited during or concomitant with the activation phase. This is in line with the general observation that partial agonists currents are strongly increased by positive allosteric modulators, indicating that the electrophysiologically silent conformations they promote are activatable and therefore not desensitized (32).

Two 5-HT-bound structures of m5-HT_3A_ displaying a closed channel sustain the intermediate reorganizations seen by VCF. Indeed, conformations called I1 (PDB 6HIO) and State 1 (PDB 6DG7) feature rearrangements at the ECD and interface between ECD/TMD similar to that of conformation displaying an open channel, but with non-conducting pore (Fig S3) (9,11). It is noteworthy that these states were initially proposed to represent either pre-active or desensitized states, and that these two possibilities could not be distinguished without ambiguity. However, indirect experiments of substituted cysteine accessibility method (SCAM) suggested that desensitization involves weak reorganizations of the upper part of the channel that holds the activation gate, arguing for the pre-active state hypothesis (9). Combined SCAM and VCF data thus provide compelling evidence for annotation as pre-active state. Interestingly, during the revision of this article, several cryo-EM structures of m5-HT_3A_ in complex with partial agonists SMP-100 and ALB-148471 were published. In full agreement with our VCF data, partial agonists are found to stabilize the protein in both active and pre-active-like conformation (32).

Intermediate conformations are also observed on many occasions for the sensors near the orthosteric site. This is the case with the var partial agonist on S204C, which produces a lower maximum current variation (ΔImax) than that caused by 5-HT but a higher ΔFmax. The phenotype is exacerbated in the presence of N101K, 5-HT and mCPBG showing intermediate conformations at sub-saturating concentrations, while varenicline promote no current while still evoking robust ΔF. Of note, on this mutant, mCPBG is the most efficient agonist, but it remains possible that it has a partial agonistic character due to the strong loss-of-function N101K mutation. In this context, the sensor I160C/Y207W shows a loss-of-function phenotype resembling that of S204C/N101K with comparable relative position of the ΔI and ΔF dose-response curves. Corelating the motions at the orthosteric site with known cryo-EM structures is, however, difficult. Indeed, two hypotheses can be sustained by these results: the intermediate conformations revealed by these sensors can correspond either to those stabilized by setrons or to the pre-active conformations discussed above.

Altogether, VCF data highlight a progressive propagation of the signal following ligand binding. Setrons elicit local reorganizations shown by the sensors located around the orthosteric site, partial agonists elicit local reorganizations at the four sensors indicating a motion of the whole ECD with partial pore opening, and strong agonists elicit reorganizations detected by all four sensors together with channel opening. VCF data thus identify four families of conformations endowed with distinct ΔI/ΔF signatures contributing to signal transduction: resting-like apo, setron-inhibited, partial agonist-elicited pre-active and active states. The topological information given by the various sensors are remarkably consistent with the gallery of high-resolution structures solved thus far. Indeed, the data from fluorescence partners are consistent with respectively Apo, setron-bound, 5-HT bound-closed and 5-HT-bound open conformations. This provides important information allowing reasonable functional annotation of the various structures to physiological states in a membrane environment, at least regarding the ECD. Figure 4 shows a speculative four-state model as a framework integrating the whole set of data. In addition, VCF data give insights in the action of partial agonists, that do not exclusively stabilize the active state and document the phenotypes of various allosteric mutations.

**Figure 4.**
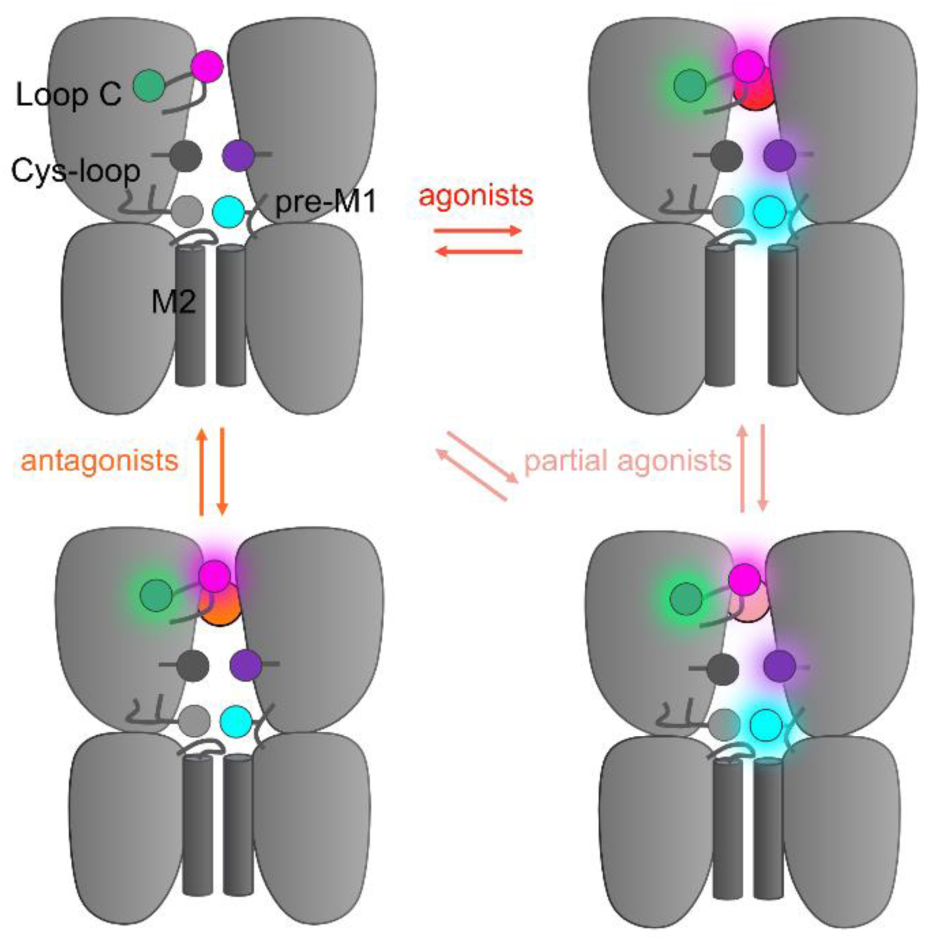
Hypothetical integrative model of VCF data. Schematic representation of the 5-HT_3_R representing two subunits in side-view with the orthosteric site and ion channel M2 helices highlighted. The four fluorescent sensors are represented as hexagons following the color code of figure 1 (S204C in magenta, I160C/Y207W in green, V106C/L131W in purple, R219C/Y140W in cyan). Ligand-elicited fluorescence changes are represented as a light halo. VCF data identifies four different conformations whose fluorescence patterns match known high-resolution structures. These conformations are called resting (matching apo structures, PDB 4PIR, 6BE1, 6H5B, 6Y59), inhibited (matching setron-bound structures, PDB 6HIS, 6W1J, 6W1M, 6W1Y, 6Y1Z), intermediate (matching I1 and state1, PDB 6HIO, 6DG7) and active (matching F, state2, and open, PDB 6HIN, 6DG8, 6Y5A).

A key observation of the study is the identification of pre-active intermediates that are favored upon binding of partial agonists and/or in the presence of loss-of-function mutations. It is noteworthy that single-channel kinetic analyses of pLGICs early showed that the activation transition pathway involves multiple intermediated states. Analysis of numerous mutants of the muscle nAChR analyzed by REFERs (rate equilibrium linear free energy relationships) suggested a multistep reorganization that initiates in the orthosteric site and progressively spreads to the ion channel gate (33). Analysis on GlyRs and nAChRs detected late intermediate-states called “flip” (34) or “primed” (35), that are favored by loss of function mutations (36,37), while a non-conducting intermediate state has been proposed from kinetic models of a high-conductance mutant of 5-HT_3_R(7).

More recently, fluorescence and VCF studies identified intermediate conformations for nAChRs, ⍺1-GlyRs and the bacterial homolog GLIC(21,38–41). Of note, pre-active intermediates were unraveled by a fluorescence quenching pair at strictly homologous positions at the ECD-TMD interface of 5-HT_3_R and ⍺1-GlyRs. Indeed, residues R219C/Y140W are homologous to another sensor pair Q219C/K143W engineered on the ⍺1-GlyR. MTS-TAMRA-labelled Q219C/K143W also reports a non-conducting intermediate in the pathway towards activation, with a fluorescence variation revealing an early reorganization of the ECD-TMD interface(21). Molecular dynamic simulations starting from a taurine-bound closed cryo-EM structure was found to recapitulate the pharmacological properties observed by VCF, suggesting that the intermediate conformations correspond to a highly dynamic family of conformations featuring an active-like ECD but a resting-like pore(42). Altogether, VCF shows that virtually all pLGICs appear to share a common global mechanism of gating where the proteins visit a family of structurally dynamic intermediates showing an active-like ECD and a resting-like TMD conformation. The present work thus extends this idea to the 5-HT_3A_R, together with providing structural blueprints for cryo-EM structural annotation.

In conclusion, by monitoring simultaneously electrophysiologically silent and active conformations of the 5-HT_3_R, VCF allowed to characterize the mechanisms of action of allosteric effectors and allosteric mutations. We found that the strength of the agonist correlated to a degree with propagation of this fluorescence change beyond the local site of neurotransmitter binding, unraveling intermediate conformations allowing proposing a structure-based functional annotation of known high-resolution structures. Besides validating the mechanism underlying inhibition by setrons, these data unravel a unique mode of action of partial agonists to promote distinct intermediate conformations. The various fluorescence sensors developed will be valuable in future mutational analysis and characterization of drugs acting at the 5-HT_3_R that hold promise for clinical use.

## Materials and Methods

### Materials

The ligands used for perfusion: 5-hydroxytryptamine hydrochloride (5-HT, serotonin), mCPBG, varenicline tartrate, alosetron, granisetron, ondansetron and metoclopramide have been purchased from Merck (Sigma). Hydrosoluble ligands were directly solubilized in distilled water or in recording solution, aliquoted at high concentrations, kept in −20°C and used freshly for dilution on the day of experiment. Less soluble ligands (ondansetron) were first dissolved in DMSO, aliquoted and kept at −20°C and then diluted in recording solutions at final concentrations not exceeding 1% of final DMSO on the day of experiment. The fluorescent dye MTS-TAMRA was purchased from Clinisciences, aliquoted in DMSO, keep at −20°C and freshly used for labelling.

### Site-directed mutagenesis

All mutations were carried into the gene of the mouse 5-HT_3A_ (described in Polovinkin et al., where strep-tags were removed), kindly provided by Hugues Nury (Institut de Biologie Structurale, Grenoble, France) (9). The mutations were introduced by site-directed mutagenesis *via* PCR (CloneAmp Hifi, Takara). All mutations were assessed by complete sequencing of the gene (Eurofins Genomics). We have chosen to work with the mouse receptor because most of the available atomic structure have been obtained for m5-HT_3A_ receptors.

### Oocytes handling

*Xenopus laevis* ovarian fragments (TEFOR PARIS SACLAY CNRS UAR2010 / INRAE UMS1451) and dissociated stage VI oocytes (Ecocyte Biosciences, Germany) were used in the context of this study. Concerning ovarian fragments, oocytes were dissociated following enzymatic treatment with collagenase II (1 mg/mL; 1h at room temperature in gentle agitation) (Thermofischer) in ORII solution (in mM: 82.5 NaCl, 2.5 KCl, 1 MgCl_2_, 5 HEPES, pH adjusted to 7.6 with NaOH). Selected oocytes were then handled in Barth’s solution (in mM: 88 NaCl, 1 KCl, 0.33 Ca(NO_3_)_2_, 0.41 CaCl_2_, 0.82 MgSO_4_, 2.4 NaHCO_3_, 10 HEPES, pH adjusted to 7.6 with NaOH) at 18°C.

### cDNA injection

cDNA encoding for m5-HT3A constructs were injected into the nucleus of oocytes (100ng) with cDNA encoding for eGFP as a reporter of correct injection (25ng) by an air injection system (Nanoject II, Drummond). Oocytes were then incubated at 18°C and used to be recorded 48 to 96 hours after injection.

### Labelling of mutants

Oocytes were incubated for 20 minutes at room temperature with a solution containing 10 μM of MTS-TAMRA (in DMSO) and 1 to 10 µM of 5-HT (in ND96 0% calcium) allowing the final concentration of DMSO to not exceed 0.1%. The oocytes were then rinsed 3 times with perfusion solution and recorded during the following two hours.

### Voltage-clamp fluorometry

Oocytes were placed in a custom-made recording chamber (described in Shi et al., 2023) (21). It allows to record and perfused the same part of the animal pole that faces the perfusion and the inverted microscope. Oocytes were continuously perfused with freshly made ND96 0% Calcium (in mM: 96 NaCl, 2 KCl, 5 HEPES, 1 MgCl2, 1 HEPES, 1 EGTA and pH was adjusted at 7.6 with NaOH). To perform recordings, micro-electrodes (Borosilicate glass with filament BF150-110-7.5, WPI) of resistances comprised between 0.2 and 2 mΩ (pipette puller PC-10, Narishige) were used and oocytes were clamped at −60 mV for all the experiments. Recordings are performed with a GeneClamp 500 voltage patch-clamp amplifier (Axon Instruments) and a 1400 A digitizer (Axon Instruments) with Clampex 10.6 software (Molecular Devices). Recorded currents were sampled at 2kHz and filtered at 500Hz. The fluorescence emission was recorded via a FF01-543_22 bandpass filter (Semrock) and collected by a photo-multiplicator (H10722, Hamamatsu). The intensity of irradiation of the LEDs (pE-4000 CoolLED) and of the sensitivity of detection of the PMT were kept at the same level for all the experiments. For the screening experiment, 5-HT at 50 and 100µM where perfused (around 10s). Dose-response curves of 5-HT, mCPBG and varenicline were adapted to the phenotype of the construct (gain or loss-of-function) and used at concentrations stated in legends of graphs from the figures. Each concentration has been repeated twice for establishment of the dose-response curves. For the pharmacological experiments, used compounds were perfused on the same oocyte at saturating concentration (5-HT and mCPBG at 200 µM, varenicline at 400 µM, antagonists used have low nanomolar affinities and used here all at 3µM concentration).

### Two electrode voltage-clamp

Currents from impaled oocytes expressing m5-HT_3A_ constructs were obtained under ND96 0% calcium perfusion. Currents were recorded by a Warner OC-725C amplifier and digitized by a Digidata 1550 A with Clampex 10 software (Molecular Devices). Currents were sampled at 500 Hz and filtered at 100 Hz. The voltage clamp is maintained at −60mV during all experiments. The TEVC setup has a faster perfusion than the VCF setup, as described in a previous paper from our lab (22). To characterize some of the desensitization properties, long application of 5-HT (45s) at several concentrations (10, 50, 100, 300 and 500 µM) have been applied to calculate the remaining currents after 45s application and compared to WT.

### Analysis of results

Current and fluorescence analyses were made with Clampfit (Molecular Devices, Sunnyvale, CA). Dose response curves, EC_50_ and Hill coefficients are obtained by the normalization of serotonin-induced currents to the maximal current followed by the fitting of the data by one-site Hill equation (GraphPad Prism). Error bars on figures represents +/− SEM. Results were obtained from at minima five different oocytes from at minima two different batches of oocytes. Rise time constants and desensitization constants have been calculated in Clampfit by mono-exponential fittings of signals. Statistical analyses have been made with Prism (unpaired t-tests).

## Acknowledgments

The work was supported by the ERC (Grant no. 788974, Dynacotine). The authors would like to thank Hugues Nury for the kind gift of the mouse 5-HT_3_ cDNA, Kate Dunning, Solène Lefebvre, and Thomas Grutter for the critical reading of the manuscript.

## Notes

### Competing Interest Statement

The authors have declared no competing interest.

### Summary of Updates

First, the manuscript has been clarified with better references to previous work. Also, to better illustrate the structural reorganizations seen in the cryo-EM structures and that are used for VCF data interpretation, we added a new supplementary figure 3. It shows a superimposition of Apo, setron and 5-HT bond structures, with reorganization of loop C and Cys-loop consistent with VCF data. We have added statistical analysis when relevant. We have also revised the figures to make them clearer and easier to read.

