## Supplementary files for "Mapping the molecular motions of 5-HT_3_ serotonin-gated channel by Voltage-Clamp Fluorometry"

**This PDF file includes:**

Figures S1 to S3
Table S1

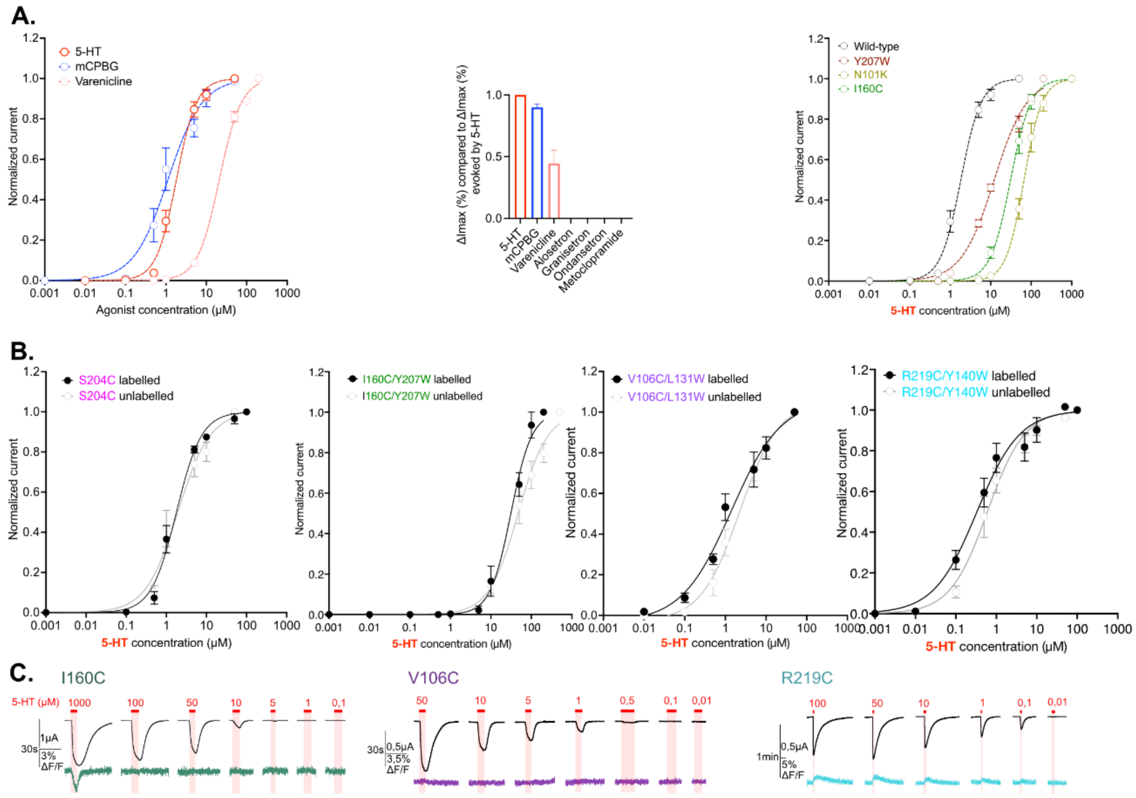

**Fig. S1. Controls. A.** Left panel: wild-type receptor currents dose-response curves for 5-HT (red), mCPBG (blue) and varenicline (salmon) on the VCF setup. Middle panel: comparison of maximal current variations evoked by saturating concentrations of 5-HT, mCPBG, varenicline, alosetron, granisetron, ondansetron and metoclopramide. Values are normalized with the maximal current variations evoked by 5-HT. Left panel: effect of each loss-of-function mutation: Y207W in burgundy, I160C in green and N101K in khaki. Dose-response curves for 5-HT obtained on the VCF setup. **B.** Characterization of the potential effect of MTS-TAMRA labelling on the four sensors developed in this study. Dose response curves of current with (black curve and dots) and without labelling (grey curve and dots) are shown. **C.** Representative recordings of I160C (left, green), V106C (middle, purple) and R219C (right, cyan) labelled with MTS-TAMRA upon different concentrations of 5-HT. Note that on the left panel (I160C), the direct fluorescence quenching evoked by high concentration of 5-HT (1000 µM) is highly visible, which is not the case at lower concentrations.

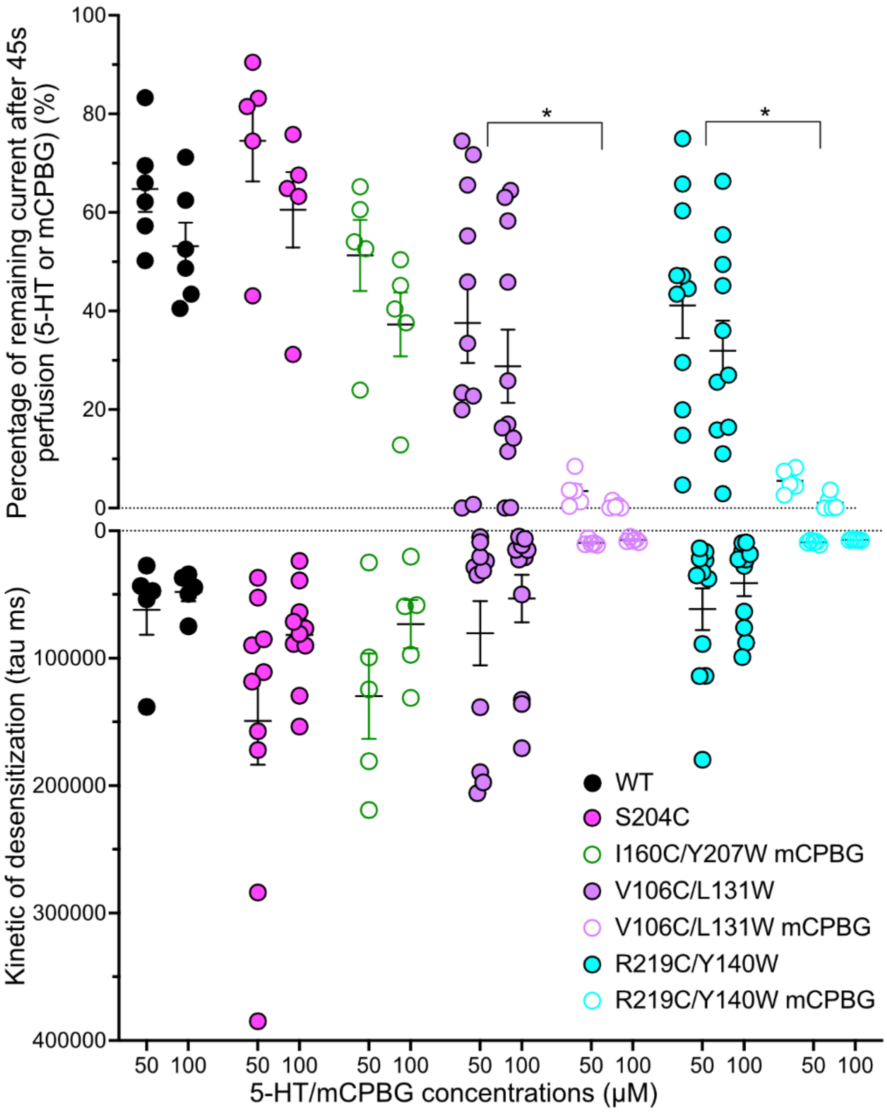

**Fig. S2. Desensitization properties**. Desensitization properties for the four sensors studied by TEVC. Top panel: graphical representation of the remaining current recorded after 45 seconds of constant perfusion of 5-HT (when nothing is mentioned) or mCPBG (see legends on the graph) at 50 and 100 µM. The remaining current (in %) is normalized to the maximum current obtained for each application. Bottom panel: graphical representation of desensitization constants tau obtained by mono-exponential fitting on the desensitization part of the recordings during long application of 5-HT or mCPBG. Note the significative difference between desensitization properties evoked by 5-HT and mCPBG on V106C/L131W and R219C/Y140W (unpaired t-tests).

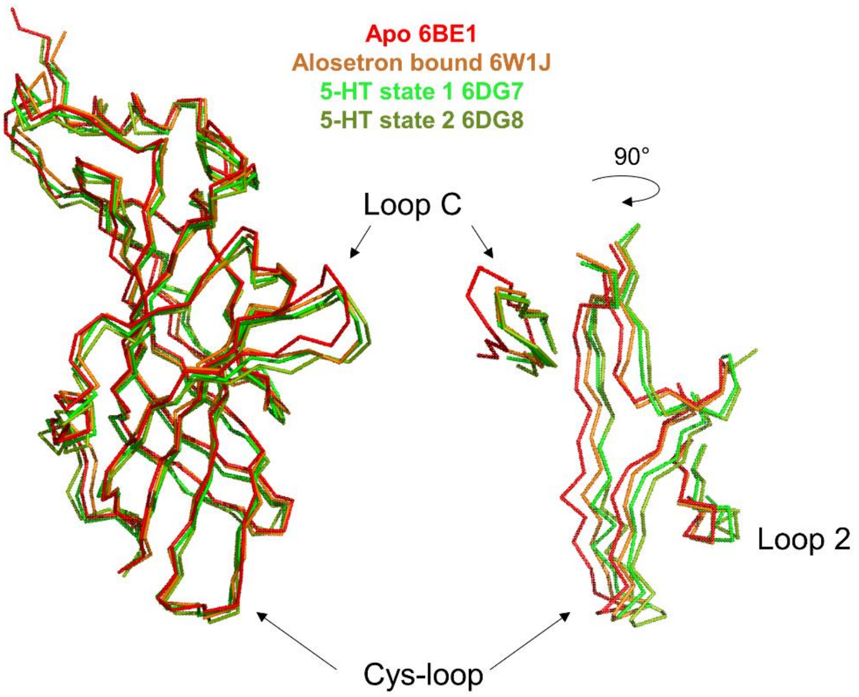

**Fig S3. Structural comparison of representative Cryo-EM structures.** Superimposition of 5-HT$_3$R structures in Apo, Alosetron-bound, 5-HT-bound state 1 (closed channel) and state 2 (open channel). On the pentamers, structures are superimposed at the level of the ECD of the subunit adjacent to the one shown (chain B) (the so-called complementary subunit facing loop-C, chain A). The superimposition highlights that Alo and 5-HT equally promote the capping of loop C, but that Alo binding is associated with weak reorganization of the lower part of the ECD especially at the level of the Cys-loop and loop 2 near the ECD-TMD interface. In contrast, both 5-HT-bound states show a similar reorganization regardless of whether the channel being open or closed.

**Table S1.** Values of $\tau_{rise}$ obtained from rise time kinetic analyses for current and fluorescence on labelled m5-$HT_{3A}$ mutants.

| Construct | Molecule | Concentrations (µM) | τ (ms) mean +/-SEM | | Unpaired t-test (p value) |
|---|---|---|---|---|---|
| S204C | 5-HT | 10 | I | 1490.106 +/- 226.23 | Ns (0.0801) |
| | | | F | 2720.210 +/- 602.472 | |
| | | 50 | I | 860.925 +/- 115.317 | Ns (0.0556) |
| | | | F | 1607.027 +/- 332.607 | |
| S204C | mCPBG | 10 | I | 780.395 +/- 68.012 | * (0.0331) |
| | | | F | 1014.284 +/- 50.880 | |
| | | 50 | I | 603.974 +/- 116.549 | Ns (0.4805) |
| | | | F | 751.475 +/- 157.765 | |
| S204C + N101K | 5-HT | 50 | I | 13744.483 +/- 1207.601 | **** (<0.0001) |
| | | | F | 2150.838 +/- 260.847 | |
| | | 100 | I | 9972.073 +/- 1247.551 | **** (<0.0001) |
| | | | F | 2040.467 +/- 177.917 | |
| S204C + N101K | mCPBG | 50 | I | 1778.377 +/- 148.304 | ** (0.0036) |
| | | | F | 1151.664 +/- 74.421 | |
| | | 100 | I | 1387.618 +/- 157.334 | * (0.0206) |
| | | | F | 919.538 +/- 65.337 | |
| I160C/Y207W | 5-HT | 50 | I | 5754.586 +/- 1010.8129 | **(0.0061) |
| | | | F | 2181.044 +/- 371.452 | |
| | | 100 | I | 5989.563 +/- 816.874 | *** (0.0008) |
| | | | F | 1828.336 +/- 202.146 | |
| I160C/Y207W | mCPBG | 10 | I | 2285.706 +/- 393.976 | Ns (0.8483) |
| | | | F | 2173.099 +/- 420.552 | |
| | | 100 | I | 921.791 +/- 0216.664 | Ns (0.0508) |
| | | | F | 2242.328 +/- 568.780 | |
| V106C/L131W | 5-HT | 1 | I | 7868.196 +/- 1860.805 | Ns (0.2294) |
| | | | F | 4928.317 +/- 1388.095 | |
| | | 10 | I | 1669.049 +/- 212.492 | Ns (0.1258) |

| | | | | |
|---|---|---|---|---|
| | | | F | 1209.388 +/- 174.794 |

| | | | | | |
|---|---|---|---|---|---|
| V106C/L131W | varenicline | 1 | I | 8099.307 +/- 2081.888 | Ns (0.0755) |
| | | | F | 2669.500 +/- 616.985 | |
| | | 100 | I | 1320.949 +/- 215.924 | Ns (0.5363) |
| | | | F | 1120.105 +/- 227.454 | |

| | | | | | |
|---|---|---|---|---|---|
| R219C/Y140W | 5-HT | 1 | I | 1884.998 +/- 323.988 | * (0.0342) |
| | | | F | 2936.035 +/- 283.908 | |
| | | 10 | I | 619 +/- 61.883 | ** (0.0080) |
| | | | F | 1490 +/- 267.086 | |
| | | 50 | I | 778.830 +/- 156.135 | ns (0.0764) |
| | | | F | 1708 +/- 453.706 | |

| | | | | | |
|---|---|---|---|---|---|
| R219C/Y140W | varenicline | 1 | I | 2417 +/- 272.127 | Ns (0.0575) |
| | | | F | 4623.726 +/- 957.392 | |
| | | 10 | I | 544.917 -+/ 32.752 | ** (0.0025) |
| | | | F | 2502.016 +/- 451.889 | |
| | | 100 | I | 397.615 +/- 59.082 | * (0.0336) |
| | | | F | 1825.950 +/- 554.506 | |

| | | | | | |
|---|---|---|---|---|---|
| R219C/Y140W + N101K | 5-HT | 1 | I | 3467.478 +/- 764.973 | Ns (0.1853) |
| | | | F | 5909.440 +/- 1536.925 | |
| | | 10 | I | 1721.749 +/- 274.859 | * (0.0111) |
| | | | F | 3199.143 +/- 433.522 | |
| | | 100 | I | 1135.772 +/- 186.094 | Ns (0.3671) |
| | | | F | 1386.235 +/- 193.816 | |

| | | | | | |
|---|---|---|---|---|---|
| R219C/Y140W + N101K | varenicline | 10 | I | 2914.184 +/- 192.065 | Ns (0.0851) |
| | | | F | 6042.400 +/- 1580.925 | |
| | | 100 | I | 2707.839 +/- 755.202 | Ns (0.5815) |
| | | | F | 3448.594 +/- 1045.416 | |
